## Appendix for "A proximity proteomics pipeline with improved reproducibility and throughput"

### Table of contents


| Content | Title | Page |
|---|---|---|
| Appendix Figure S1 | Schematic of compartment-specific APEX2-tagged constructs | 3 |
| Appendix Figure S2 | Sample enrichment and DIA-mass spectrometry method optimization. | 4 |
| Appendix Figure S3 | Comparison of manual- and automated- enrichment methods coupled to DDA- and DIA-mass spectrometry methods. | 5 |
| Appendix Figure S4 | Characterization of all selected monoclonal APEX2-tagged localization domain cell lines. | 6 |
| Appendix Figure S5 | APEX-labeling proteomics quality control. | 7 |
| Appendix Figure S6 | Volcano plots for APEX2 proximity labeling of each subcellular compartment. | 8 |
| Appendix Figure S7 | Mapping ligand-dependent proximal interaction network changes of $5HT_{2A}$. | 9 |
| Appendix Figure S8 | $5HT_{2A}$ network dynamics for sustained, activity-dependent proximal interactions. | 10 |
| Appendix Figure S9 | Flow cytometric analysis of $5HT_{2A}$ receptor at the plasma membrane. | 11 |
| Table EV1 | Performance comparison of the automated PL strategy combined with DIA-based MS. | 12 |
| Table EV2 | Summary of optimized conditions in high-input vs. low-input proximity proteome pipeline. | 13 |

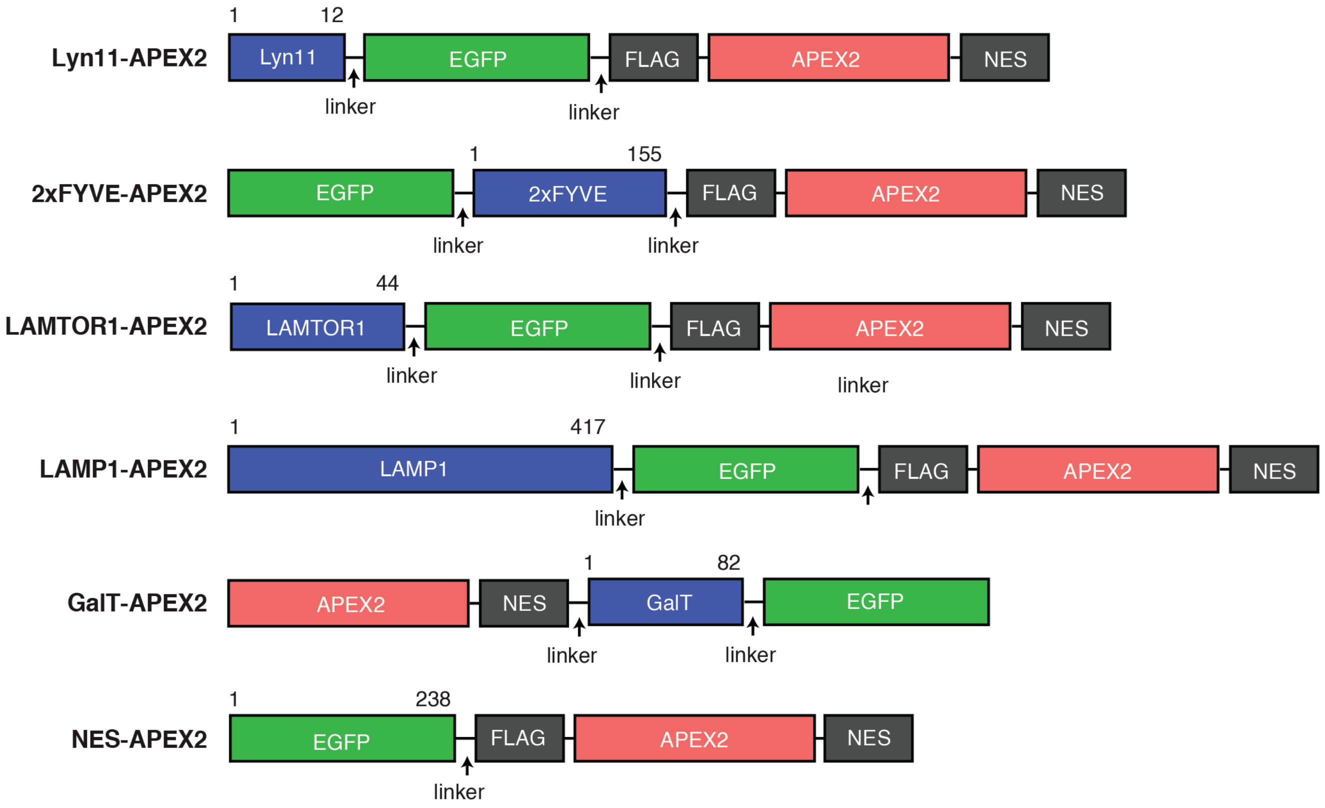


***Appendix Figure S1. Schematic of compartment-specific APEX2-tagged constructs.***

In our study, we selected protein localization domain Lyn11 for plasma membrane, 2xFYVE for endosome, LAMTOR1 and LAMP1 targeting sequence for late endosome/lysosome, and β-1,4 galactosyltransferase (GalT) for Golgi apparatus. A non-location specific construct for the cytosol was used as a control.

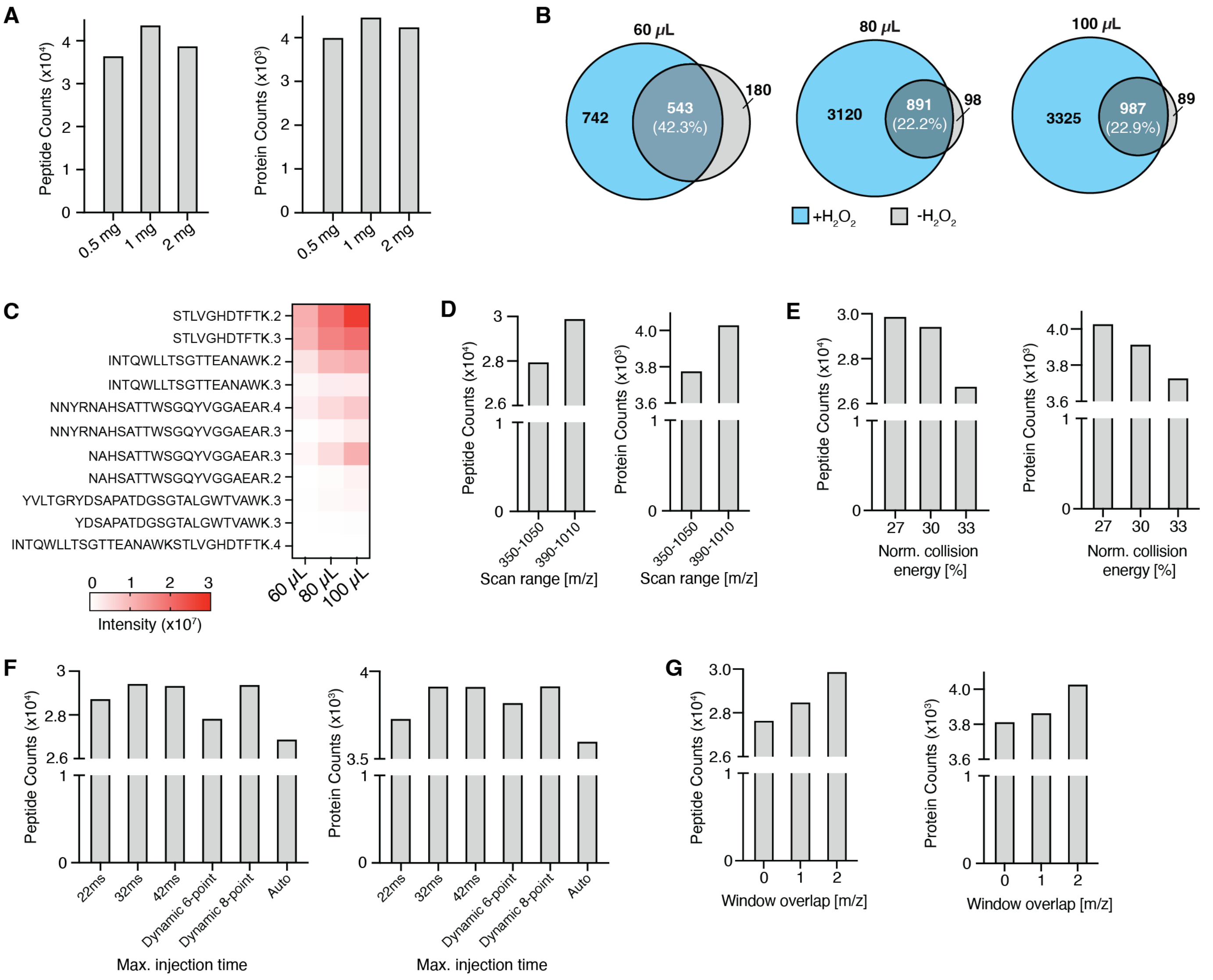


***Appendix Figure S2. Sample enrichment and DIA-mass spectrometry method optimization.***

**A**, Different protein loading amounts with 100 μL beads. **B**, Venn diagram of proteins identified with vs. without $H_2O_2$ treatment using different beads. **C**, Line chart of streptavidin peptides intensities. Three replicates for each condition. **D-G**, Optimization of DIA-mass spectrometry parameters including: Scan range (**D**), Normalized collision energy (**E**), Maximum injection time (**F**), Width of overlapping window (**G**).

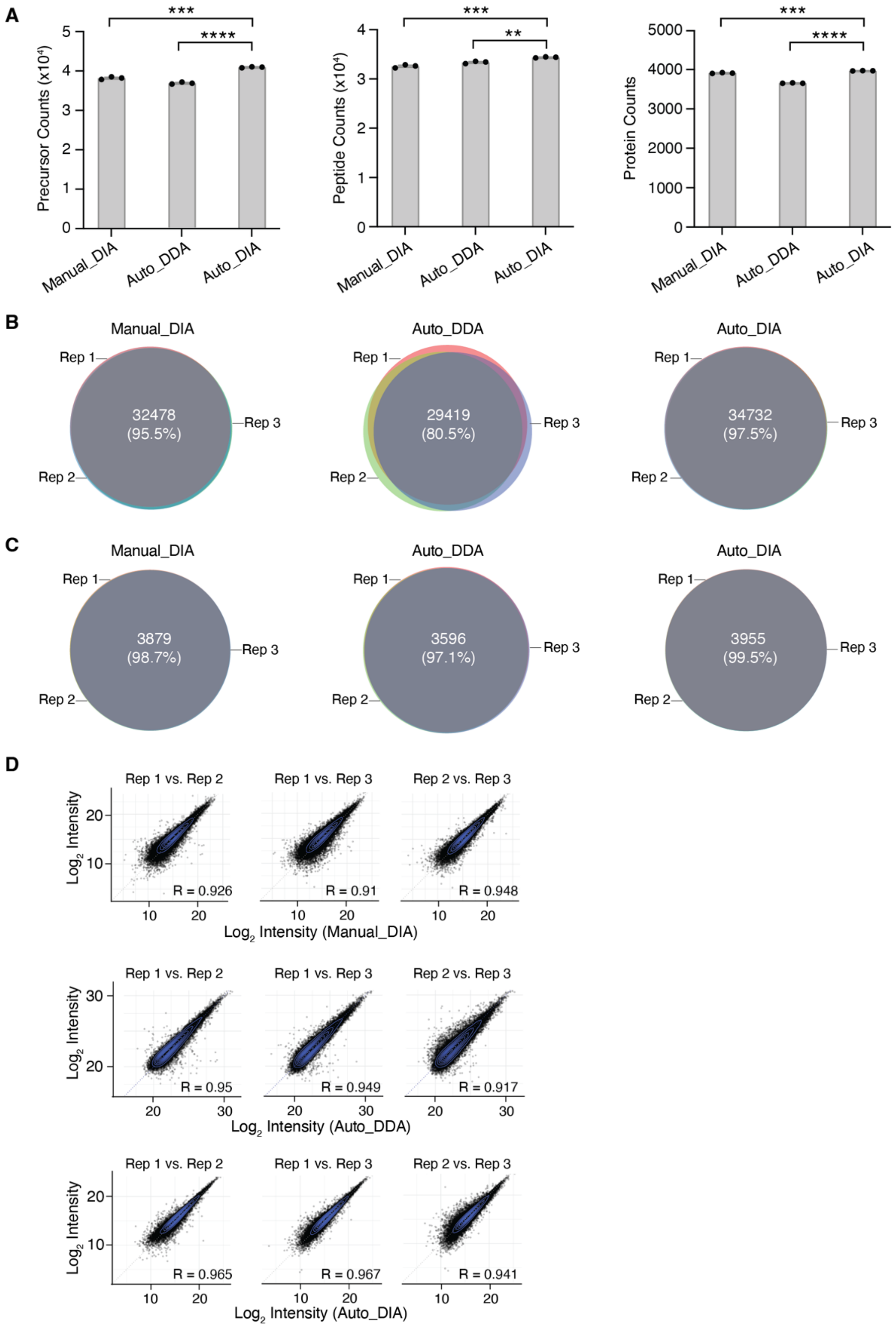


***Appendix Figure S3. Comparison of manual- and automated- enrichment methods coupled to DDA- and DIA-mass spectrometry methods.***

**A**, Number of precursors, peptides, and proteins being identified in the three methods. Statistical significance was performed in Prism (GraphPad) using unpaired t-test. **B**, Venn diagram analysis comparing peptide identification across three replicates. **C**, Venn diagram analysis comparing protein identification across three replicates. **D**, Correlation analysis of peptide intensities.

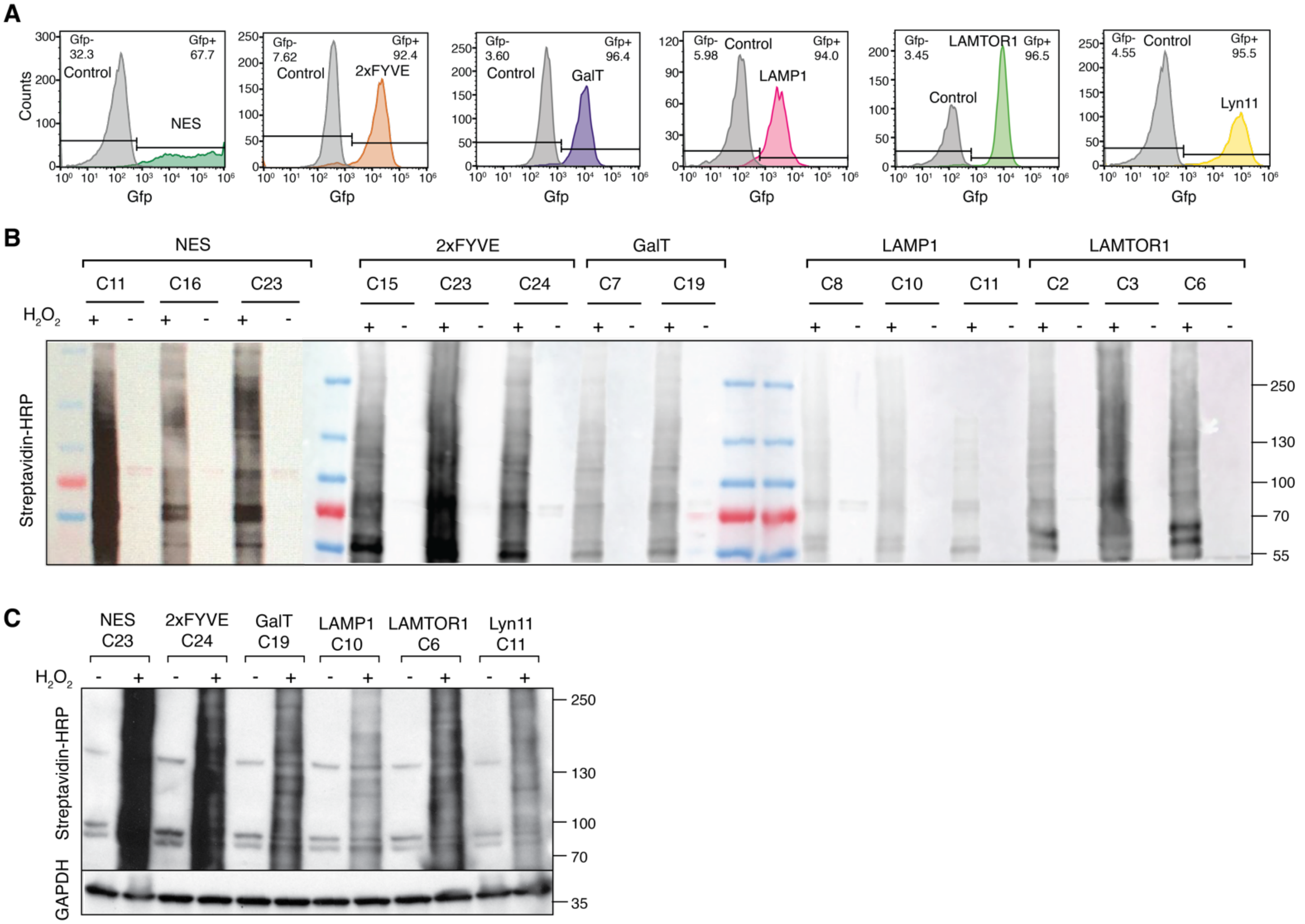


***Appendix Figure S4. Characterization of all selected monoclonal APEX2-tagged localization domain cell lines.***

**A**, Flow cytometry analysis of APEX2-tagged location domain cell lines. GFP was used to evaluate expression and localization of the APEX2 construct. The non-Doxycycline induced cell line was used as control. **B**, Western blot analysis of whole cell lysate derived from monoclonal APEX2 cell lines. **C**, Western blot analysis of final monoclonal APEX2-tagged location domain cell lines with or without $H_2O_2$.

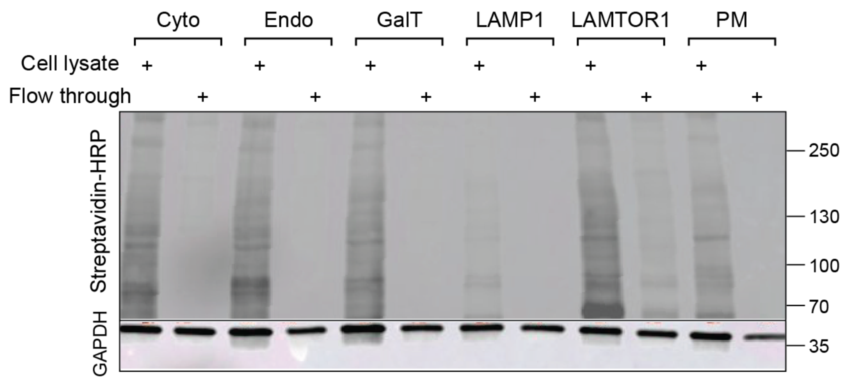


***Appendix Figure S5. APEX-labeling proteomics quality control.***

Western blot analysis of biotinylation and enrichment efficiency of 6 monoclonal APEX2-tagged location domain cell lines.

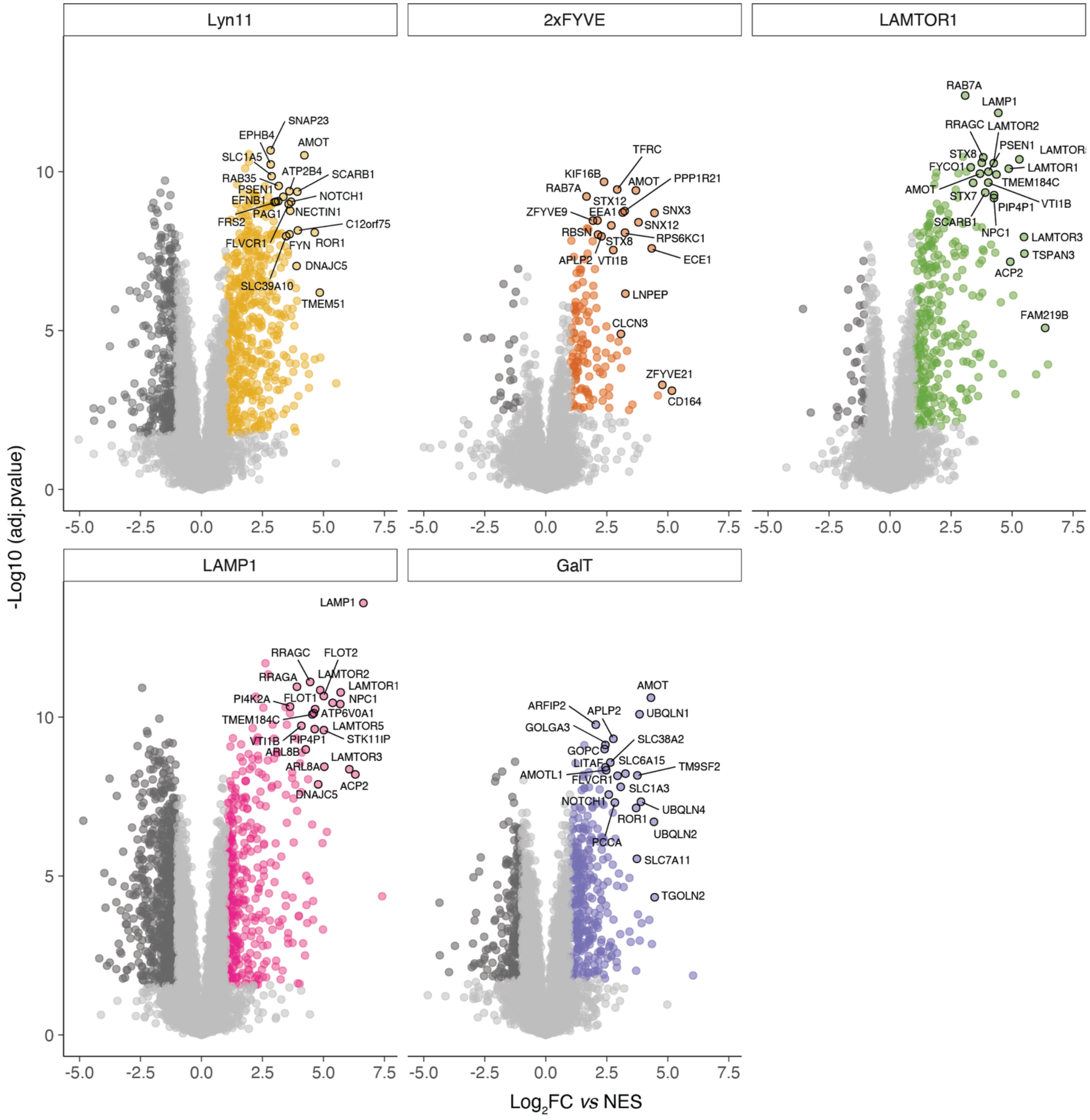


***Appendix Figure S6. Volcano plots for APEX2 proximity labeling of each subcellular compartment.*** Enlarged volcano plots of **Figure 4C** to annotate known location specific proteins. Proteins with a $log_2$ fold change >1 and adjusted p-value <0.05 were considered significant and colored.

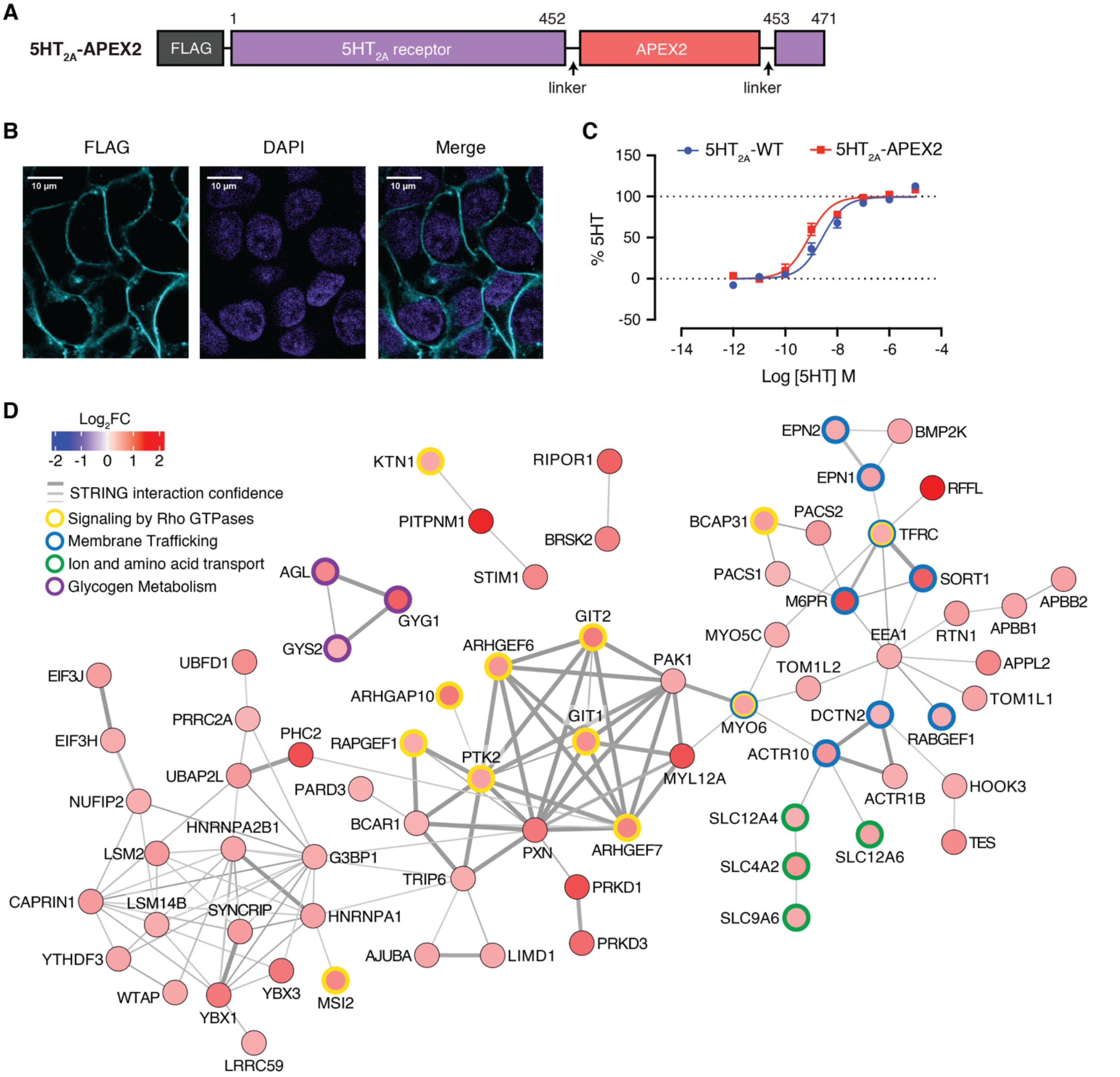


***Appendix Figure S7. Mapping ligand-dependent proximal interaction network changes of 5HT$_{2A}$.***

**A**, Construct design of 5HT$_{2A}$ receptor. **B**, Confocal imaging of monoclonal 5HT$_{2A}$-APEX2 cell lines. Cells were stained with DAPI for the nucleus and the APEX2 construct location was indicated by FLAG. Scale bar represents 10 µm. **C**, BRET validation of 5HT mediated Gq recruitment to APEX2-tagged vs. wild-type 5HT$_{2A}$. **D**, Protein interaction network connecting proteins with sustained agonist-dependent changes in the proximity of 5HT$_{2A}$. Proteins shown as nodes were colored according to their $\log_2$ fold change. The edges connecting the proteins were derived from STRING (Szklarczyk *et al*, 2011) and the edge width was scaled according to the interaction confidence. Proteins corresponding to Reactome pathways (Jassal *et al*, 2020) that were enriched within the sustained cluster were indicated with different colored node borders.

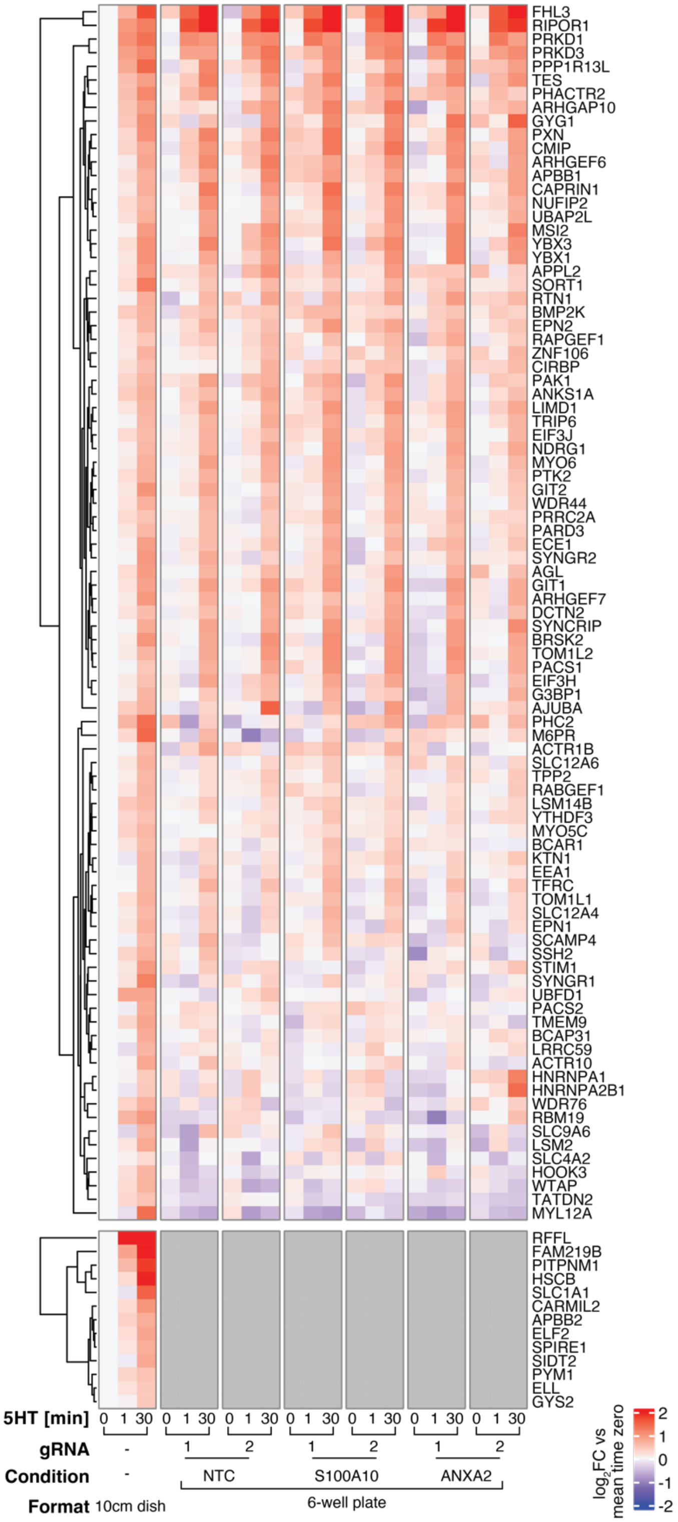


***Appendix Figure S8. 5HT$_{2A}$ network dynamics for sustained, activity-dependent proximal interactions.***

Heatmap depicting proteins from 5HT2A-APEX2 experiment with sustained responses to 5HT treatment (**Appendix Figure S7D**) compared across NTCs and knockouts of S100A10 and ANXA2 in 6-well plate format and the 5HT2A APEX2 data from 10cm dishes. Data were collected from three independent biological replicates (n = 3) for NTCs and S100A10 and two independent biological replicates (n = 2) for ANXA2.

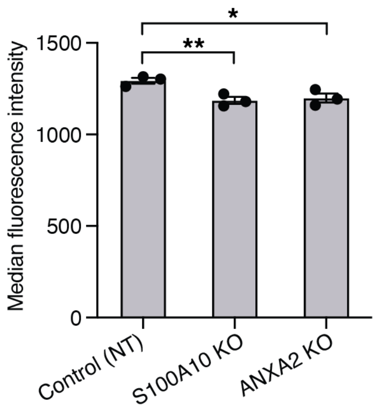


***Appendix Figure S9. Flow cytometric analysis of $5HT_{2A}$ receptor at the plasma membrane.***

Quantification of $5HT_{2A}$ receptor at the plasma membrane in NTC, S100A10 KO, and ANXA2 KO cells. Data from three independent experiments are presented as mean ± SEM. Statistical significance was performed in Prism (GraphPad) using unpaired t-test (NTC vs. S100A10 KO, *p* value: 0.006; NTC vs. ANXA2 KO, *p* value: 0.0161.)

| | Rep 1 total | Rep 2 total | Rep 3 total | Rep 1 only | Rep 2 only | Rep 3 only | Rep 1 ∩ Rep 2 | Rep 2 ∩ Rep 3 | Rep 3 ∩ Rep 1 | Rep 1 ∩ Rep 2 ∩ Rep 3 |
|---|---|---|---|---|---|---|---|---|---|---|
| Manual DIA | 37867 | 38502 | 38216 | 163 | 49 | 55 | 502 | 959 | 210 | 36992 (95%) |
| Auto DDA | 36875 | 36618 | 36408 | 604 | 946 | 954 | 2373 | 1556 | 2155 | 31743 (78.7%) |
| Auto DIA | 41058 | 40991 | 40796 | 45 | 74 | 55 | 489 | 217 | 313 | 40211 (97.1%) |

***Table EV1. Performance comparison of the automated PL strategy combined with DIA-based MS.*** Details for the Venn diagrams of precursor features depicted in **Figure 3E**.

| | High-input PL | Low-input PL |
|---|---|---|
| Cell culture | 10cm dish | 6-well plate coated with Poly-D-lysine (PDL) |
| Cell seeding density | 4 x $10^6$ cells/dish | 0.5 x $10^6$ cells/well |
| Sample input amount | 1 mg | 0.25 mg |
| Beads amount | 80 µL | 25 µL |
| Enrichment buffer volume | 1 mL | 200 µL |
| Beads washing volume | 1 mL | 200 µL |
| Protein digestion volume | 200 µL | 100 µL |
| MS sample loading amount | Resuspended in 20 µL 0.1% formic acid and inject 1 µL | Resuspended in 20 µL 0.1% formic acid and inject 3 µL |

***Table EV2. Summary of optimized conditions in high-input vs. low-input proximity proteome pipeline.***
