## Supplementary material for "A proximity proteomics pipeline with improved reproducibility and throughput": Dataset EV2

### Steps data

| | | |
|---|---|---|
| Tip1 | 96 DW tip comb | |
| Pick-Up | Tips | |
| 1st Beads Wash | Bead Wash 1 | |
| Beginning of step | Precollect | No |
| | Release beads | No |
| Mixing / heating: | Shake 1 time, speed | 00:00:02, Half mix |
| | Shake 2 time, speed | 00:00:30, Slow |
| | Heating during mixing | No |
| End of step | Postmix | No |
| | Collect count | 5 |
| | Collect time [s] | 10 |
| 2nd Beads Wash | Bead Wash 2 | |
| Beginning of step | Precollect | No |
| | Release beads | Yes |
| Mixing / heating: | Mixing time, speed | 00:00:30, Slow |
| | Heating during mixing | No |
| End of step | Postmix | No |
| | Collect count | 5 |
| | Collect time [s] | 10 |
| Sample Binding | Sample Binding | |
| Beginning of step | Precollect | No |
| | Release time, speed | 00:00:30, Medium |
| Mixing / heating: | Shake 1 time, speed | 06:00:00, Slow |
| | Shake 2 time, speed | 00:05:00, Half mix |
| | Shake 3 time, speed | 06:00:00, Slow |
| | Loop count | 3 |
| | Heating during mixing | No |
| End of step | Postmix | No |
| | Collect beads | No |
| Leave | Sample Binding | |

### Steps data

| | | |
|---|---|---|
| **Tip1** | **96 DW tip comb** | |
| **Pick-Up** | **Sample Binding** | |
| **Sample recollection** | **Sample Binding** | |
| Beginning of step | Precollect | No |
| | Release beads | No |
| Mixing / heating: | Mixing time, speed | 00:00:15, Half mix |
| | Heating during mixing | No |
| End of step | Postmix | No |
| | Collect count | 5 |
| | Collect time [s] | 10 |
| **1st RIPA Wash** | **RIPA Wash 1** | |
| Beginning of step | Precollect | No |
| | Release time, speed | 00:00:10, Medium |
| Mixing / heating: | Mixing time, speed | 00:03:00, Slow |
| | Heating during mixing | No |
| End of step | Postmix | No |
| | Collect count | 5 |
| | Collect time [s] | 10 |
| **2nd RIPA Wash** | **RIPA Wash 2** | |
| Beginning of step | Precollect | No |
| | Release time, speed | 00:00:10, Medium |
| Mixing / heating: | Mixing time, speed | 00:03:00, Slow |
| | Heating during mixing | No |
| End of step | Postmix | No |
| | Collect count | 5 |
| | Collect time [s] | 10 |
| **3rd RIPA Wash** | **RIPA Wash 3** | |
| Beginning of step | Precollect | No |
| | Release time, speed | 00:00:10, Medium |
| Mixing / heating: | Mixing time, speed | 00:03:00, Slow |
| | Heating during mixing | No |
| End of step | Postmix | No |
| | Collect count | 5 |
| | Collect time [s] | 10 |
| **KCL** | **1M KCl Wash** | |
| Beginning of step | Precollect | No |
| | Release time, speed | 00:00:30, Medium |
| | Mixing | [none] |
| | Heating during mixing | No |
| End of step | Postmix | No |
| | Collect count | 5 |
| | Collect time [s] | 10 |

| | | | |
|---|---|---|---|
| | Na2CO3 | Sodim carbonate wash | |
| | Beginning of step | Precollect | No |
| | | Release time, speed | 00:00:30, Medium |
| | | Mixing | [none] |
| | | Heating during mixing | No |
| | End of step | Postmix | No |
| | | Collect count | 5 |
| | | Collect time [s] | 10 |
| | Urea | 2M urea wash | |
| | Beginning of step | Precollect | No |
| | | Release time, speed | 00:00:30, Medium |
| | | Mixing | [none] |
| | | Heating during mixing | No |
| | End of step | Postmix | No |
| | | Collect count | 5 |
| | | Collect time [s] | 10 |
| | 1st Tris Wash | Tris Wash 1 | |
| | Beginning of step | Precollect | No |
| | | Release time, speed | 00:00:30, Medium |
| | Mixing / heating: | Mixing time, speed | 00:01:00, Half mix |
| | | Heating during mixing | No |
| | End of step | Postmix | No |
| | | Collect count | 5 |
| | | Collect time [s] | 10 |
| | 2nd Tris Wash | Tris Wash 2 | |
| | Beginning of step | Precollect | No |
| | | Release time, speed | 00:00:30, Medium |
| | Mixing / heating: | Mixing time, speed | 00:01:00, Half mix |
| | | Heating during mixing | No |
| | End of step | Postmix | No |
| | | Collect count | 5 |
| | | Collect time [s] | 10 |
| | Release Beads | Digestion buffer | |
| | | Release time, speed | 00:02:00, Half mix |
| | Leave | Digestion buffer | |
